# *K*-mer Genome-wide Association Study for Anthracnose and *BCMV* Resistance in the Andean Diversity Panel

**DOI:** 10.1101/2024.07.29.605481

**Authors:** Andrew T. Wiersma, John P. Hamilton, Brieanne Vaillancourt, Julia Brose, Halima E. Awale, Evan M. Wright, James D. Kelly, C. Robin Buell

## Abstract

Limited genomic resources and closely linked marker-trait associations for common beans (*Phaseolus vulgaris* L.) have limited breeders from fully utilizing molecular genetics technologies to maximize genetic gain. The emergence of virulent races of anthracnose (caused by *Colletotrichum lindemuthianum*) and *Bean Common Mosaic Virus* (BCMV) highlight the need for improved methods to identify and incorporate pan-genomic variation in breeding for disease resistance. We sequenced the *P. vulgaris* Andean Diversity Panel (ADP) and performed a genome-wide association study (GWAS) to identify associations for resistance to BCMV and eight races of anthracnose. Historical single nucleotide polymorphism (SNP)-chip and phenotypic data enabled a three-way comparison between SNP-chip, reference-based whole genome shotgun sequence (WGS)-SNP, and reference-free *k*-mer GWAS. Across all traits, there was excellent concordance between SNP-chip, WGS-SNP, and *k*-mer GWAS results—albeit at a much higher marker resolution for the WGS data sets. Significant *k-mer* haplotype variation revealed selection of the linked *I*-gene and *Co-u* traits in North American breeding lines and cultivars. Due to *k*-mer mapping criteria and the absence of target loci in the reference genome due to structural variation, only 9.1 to 47.3% of the significantly associated *k*-mers were mapped to the reference genome. To determine the genetic context of *cis*-associated *k*-mers, we generated whole genome assemblies of four ADP accessions and identified an expanded local repertoire of disease resistance genes associated with resistance to anthracnose and BCMV. With access to variant data in the context of a pan-genome, high resolution mapping of agronomic traits for common bean is now feasible.

**CORE IDEAS:**

- *K*-mer-based GWAS offers new advantages for mapping pan-genomic variation
- Comparison of reference-based SNP to reference-free *k*-mer GWAS
- Novel discovery of *cis*-associated *k*-mers for dry bean disease resistance

**PLAIN LANGUAGE SUMMARY:** Improving disease resistance in crop species such as bean is critical. We surveyed the genomes of a diverse set of bean lines and identified sequences associated with resistance to a fungal and viral pathogen. Access to the genomes of this diversity panel of beans will permit additional discoveries on the role of structural variation in phenotypes, including disease resistance.

## 1 INTRODUCTION

Crop health and food security are vulnerable to virulent plant pathogens, and improved methods are needed to identify disease resistance genes more efficiently and accurately. Common bean (*Phaseolus vulgaris* L.) is an important food crop in the US and worldwide (Uebersax et al. 2022), and includes both garden beans (green, French, string, snap beans) and dry beans (black, navy, great northern, pinto, kidney beans, etc.). Beans are an excellent source of protein, complex carbohydrates, dietary fiber, vitamins, and minerals (Siddiq & Uebersax 2022). In recent years, between 500-800,000 hectares of dry edible beans were harvested in the United States, and in 2023, a total of 1.14 million metric tons of dry beans were produced (USDA-NASS). Among the many diseases impacting common bean, anthracnose (caused by *Colletotrichum lindemuthianum*) and *Bean Common Mosaic Virus* (BCMV) are among the most devastating diseases and can cause 100% and 70% yield losses, respectively (Graham and Ranalli 1997, Hampton 1975). Anthracnose and BCMV are also seed-borne diseases and particularly problematic because they can spread rapidly through distribution and propagation of contaminated seed. Despite ongoing efforts to identify novel sources of resistance and map resistance genes and quantitative trait loci (QTL), major-gene disease resistance continues to breakdown as more virulent races of the pathogen emerge (Kelly et al. 1994, 2021). As a result, farmers must resort to more intensive management practices to suppress disease, and the seed industry must rely on strict quarantine laws and field inspections to prevent disease contaminated seed lots. A more sustainable solution is the development of crop varieties with durable genetic resistance—such as quantitative disease resistance or resistance gene pyramids.

Map-based cloning of resistance genes is difficult, labor intensive, and continues to impede efforts to breed crops with durable disease resistance. Most cloned plant resistance genes belong to two primary gene families, nucleotide binding site leucine rich repeat proteins (NLRs, 61%) and receptor-like proteins or kinases (19%), that perceive pathogen invasion directly or indirectly and trigger plant immune response (Kourelis and van der Hoom 2018). Considering the vast diversity of resistance gene homologs in the plant kingdom, relatively few resistance genes have been cloned. Moreover, of the 314 resistance genes that have been cloned, only 128 have a suggested mechanism (Kourelis and van der Hoom 2018). Key challenges that complicate positional cloning include low recombination rate at the target locus, the necessity for large segregating populations to achieve high genomic mapping resolution, a single reference genome that does not adequately represent species-wide genetic diversity, and large, heterozygous, and/or polyploid genomes. Consequently, resistance gene mapping by linkage, QTL, or a genome-wide association study (GWAS) leads to identification of genomic intervals harboring multiple candidate genes and linked DNA polymorphisms—but not necessarily the causal gene. The fact that resistance genes often occur in clusters with structural variation among accessions further exacerbates the issue. While closely linked markers can be used to select for individuals with desirable traits or haplotypes, recombination can break linkage and linkage drag can impede genetic gain.

Novel approaches are needed for rapid discovery of functional resistance gene candidates. To overcome obstacles limiting positional cloning of resistance genes, new approaches have been developed. Some of the most successful methods include MutMap (Abe et al. 2012, Takagi et al. 2013), MutChromSeq (Sanchez-Martin et al. 2016), MutRenSeq (Steuernagel et al. 2016, Witek et al. 2016), cultivar-specific long-range chromosome assembly (Thind et al. 2017), and whole genome resequencing of resistant and susceptible bulks (Wang et al. 2018). In general, each new approach combines high throughput DNA sequencing (long and short read platforms) with genomic or bioinformatic complexity reduction methods such as bulked segregant analysis, long-range linkage assembly, chromosome flow sorting, transcriptomics, and exome capture including resistance gene enrichment sequencing (Andolfo et al. 2014, Periyannan 2017). A few of these methods also rely on the development of susceptible mutant genotypes to map the resistance locus or to narrow down the list of candidate genes with independent induced mutations.

In 2018, Arora et al. demonstrated, a novel and rapid approach to clone multiple resistance genes simultaneously. By leveraging association genetics and resistance gene enrichment sequencing, researchers cloned four stem rust resistance genes from a diverse population of *Aegilops tauschii* accessions (a wild relative of bread wheat). The method relied on pangenome variation for resistance gene sequences across 174 *Ae. tauschii* accessions and differential disease response to seven *Puccinia graminis* f. sp. *tritici* races. Three fundamental advantages of the approach were 1) it by-passed the development of biparental mapping or mutagenesis populations, 2) it used *k*-mer based associations to identify sub-sequences that have diverged from the reference, and 3) the sequence dataset could be reused and augmented to increase statistical power or identify additional marker-trait-associations with new phenotypic datasets (Arora et al. 2018).

In 2020, another advancement was made by Voichek and Weigel when they demonstrated reference-free GWAS based on *k*-mers (Voichek and Weigel 2020). Unlike Arora et al. which relied on resistance-gene enrichment sequencing (i.e., a small subset of the whole genome), Voichek and Weigel utilized *k*-mers from whole genome shotgun sequence (WGS) data for *Arabidopsis thaliana*, tomato, and maize. By using WGS data, the outcome of GWAS was not restricted to sequences resembling resistance genes predominantly NLRs. This new method also reversed the traditional GWAS sequence of steps. Rather than beginning with a set of variants with known genomic locations, they identified marker-trait-associations first and subsequently identified the genomic context of those variants. In this way, GWAS was no longer reliant on the availability of a reference genome. In addition to making GWAS more accessible to crops without a reference genome, it also addressed some biases inherent to SNP-based GWAS. Unlike SNPs, *k*-mers can also be used to mark structural variants like copy number variation, presence-absence variation, insertions/deletions, and translocations/inversions (Gupta 2021a). Lastly, while SNPs typically have a defined location in the reference genome, *k*-mers can remain unanchored and enable identification of *cis*-associated *k*-mers that are absent from the reference genome but present in the pangenome of the species (Gaurav et al. 2022).

These recent advances in genome wide association analyses also demonstrate a broader transition from a single-genome approach to a pan-genome approach (Gupta 2021b). In several crop species, *de novo* assembly of multiple accessions has become necessary to represent a larger proportion of the genetic diversity present within the species (Gage et al. 2019, Walkowiak et al. 2020). As new genome assemblies and increasingly large genomic datasets are produced, plant breeders need improved bioinformatic tools (such as *k*-mer GWAS) to effectively utilize pan_-_genomic data for crop improvement (Gaurav et al. 2022, Kim et al. 2020).

Access to multiple reference genomes is common in a subset of crop species, including common bean. Due to the parallel and independent domestication of common beans in Mesoamerica and South America, there are two distinct gene pools (Schmutz et al. 2014). Beans belonging to the Andean gene pool are generally larger and include the kidney and cranberry bean market classes in the U.S., while Mesoamerican beans include small- to medium-sized black, navy, pink, small red, great northern, and pinto beans (among several other globally recognized market classes). The first reference genome for common bean was the inbred landrace line G19833 that belongs to the Andean pool (Schmutz et al. 2014). In 2016, the genotype BAT93 was sequenced to represent the genetic diversity of beans from the Mesoamerican gene pool (Vlasova et al. 2016). In addition to the *de novo* genome assembly of the G19833 and BAT93, a panel of 37 varieties were sequenced by an international collaboration to identify inter-gene pool introgressions (Lobaton et al. 2018), and 683 common bean accessions were sequenced to identify yield component traits across a north-south cline in primarily Chinese breeding germplasm (Wu et al. 2020).

The objectives of this study were three-fold: 1) develop a comprehensive genomic resource for beans from the Andean gene pool, 2) validate SNP and *k*-mer-based GWAS results, and 3) identify *k*-mers associated with disease resistance phenotypes. The Andean Diversity Panel (ADP, Cichy et al. 2015) has been widely used in the community with publicly available SNP-chip genotype as well as a broad spectrum of phenotype data including race-specific anthracnose data. Using the ADP, we performed a direct three-way comparison between GWAS results using SNP-chip data (Zuiderveen et al. 2016), WGS SNP data, and *k*-mer data. Due to the higher marker resolution and the advantages afforded from a *k*-mer-based approach, novel *cis*- associated *k*-mers could be identified, independent of a reference genome, for both anthracnose and BCMV.

## 2 MATERIALS AND METHODS

### 2.1 Plant materials and phenotypic evaluation

The ADP is comprised of large-seeded bean varieties and landraces collected from Africa, Asia, Europe, and the Americas (Song et al. 2015, http://arsftfbean.uprm.edu/bean/). In this study, a total of 241 bean lines were used, of which, 227 lines are a subset of the ADP, and 14 were included as controls (Supplemental Table S1). Twelve of the controls make up the Anthracnose differential series (Padder et al. 2017), and the remaining two controls are near_-_isogenic lines of the commercial variety cv. ‘Adams’ which are homozygous for *Co-4^2^* and *co_-_4^2^*, respectively (Kelly et al. 2021). The panel of lines used for this study closely resembles the panel evaluated by Zuiderveen et al. (2016).

Plant determinacy data from Cichy et al. (2015) was validated under greenhouse conditions at Michigan State University (14 h day length, 20°C) and scored as a case-control study where ‘1’ was used for determinate plant types and ‘0’ was used for indeterminate plant types. Bean Common Mosaic Necrosis Virus (BCMNV) phenotypic data was available online (http://arsftfbean.uprm.edu/bean/) and was coded as a case-control where ‘1’ was used for plants exhibiting a susceptible *i*-gene phenotype and ‘0’ for plants exhibiting the unique systemic necrosis *I*-gene phenotype (Bello et al. 2014, Meziadi et al. 2015, Meziadi et al. 2017). Anthracnose phenotypic data was obtained from Zuiderveen et al. (2016). Response to anthracnose was rated on a scale of 0 to 5 (described previously) where resistant plants that had no response were rated ‘0’ and plants that were highly susceptible were rated ‘5’. The mean anthracnose response of six replicates was used for association mapping. All phenotypic data used in this study are available in Supplemental Table S1.

### 2.2 Andean Diversity Panel DNA isolation, library preparation, and Illumina sequencing

Following germination, leaf tissue was collected from seedlings using forceps. A single newly emerged leaflet, approximately 1 cm in length, was collected into a tube equipped with a stainless-steel ball bearing. All tissue was collected directly into 96-well plates, with a predetermined plate layout. Following lyophilization of leaf tissue, automated tissue grinding was done using an oscillating tabletop mill (Mixer Mill MM 400, Retsch, Haan, Germany). DNA was isolated using the Mag-Bind® Plant DNA Plus 96 Kit (Omega Bio-Tek, Inc. Norcross, GA) and an automated purification system (Thermo Scientific™ KingFisher™ Flex Purification System, Waltham, MA). RNase A was used to eliminate potential RNA contamination. DNA concentration was determined using Qubit™ dsDNA High Sensitivity Assay Kit (Thermo Fisher Scientific, Waltham, MA), according to the manufacturers protocol.

DNA library preparation was done according to the manufacturers protocol using the KAPA HyperPlus Kit (Roche, Basel, Switzerland). All library preparation was done using half of the recommended volumes. A total of 200 ng of input DNA was used for each library, and the incubation period for enzymatic fragmentation was set to 7 minutes for optimal fragment size distribution. Following end repair and A-tailing, and subsequent adapter ligation, a double-sided size selection step was added to the post-ligation cleanup to select the desired library range (300 to 750 bp range). KAPA Pure Beads (Roche, Basel, Switzerland) were used for double-sided size selection at 0.5x and 0.7x concentrations. For downstream sample pooling, the KAPA Unique Dual-Indexed Adapter Kit (Roche, Basel, Switzerland) was used to index libraries with 96 unique-dual indices. Libraries were amplified using the following protocol: 1 cycle of initial denaturation at 98°C for 45 seconds; 3 cycles of 98°C for 15 seconds, 60°C for 30 seconds, and 72°C for 30 seconds; 1 cycle of final extension at 72°C for 1 minute. Lastly, 1x post_-_amplification cleanup was done using KAPA Pure Beads, and libraries eluted into 15 μl of elusion buffer. The final DNA library concentration was determined using a Qubit™ dsDNA High Sensitivity Assay Kit (Thermo Fisher Scientific, Waltham, MA), according to the manufacturers protocol.

Sequencing was done at the MSU Research Technology Support Facility using the Illumina HiSeq 4000 platform (150 nt paired end reads). A total of 90 ng of each DNA library was pooled for sequencing (pools of 15 to 30 samples) and sequenced on a single lane. Pooled samples were quantified using a Qubit™ dsDNA High Sensitivity Assay Kit (Thermo Fisher Scientific, Waltham, MA), and library size distribution was determined using an Agilent 4200 TapeStation D1000 HS (Agilent Technologies Inc., Waldbronn, Germany). The average size of pooled libraries was 493 bp. After subsampling reads from three samples with exceptionally high read counts, the average whole genome sequence coverage across all samples was 11.85x coverage, with a minimum of 8.34x and a maximum of 16.55x coverage (based on a *P. vulgaris* genome size of 587-Mb; Supplemental Table S2; Schmutz et al. 2014). Sequence quality metrics were checked using FastQC v. 0.11.8 (https://www.bioinformatics.babraham.ac.uk/projects/fastqc/) and MultiQC v. 1.0 (Ewels et al., 2016). Reference genome sequence reads, from the reference genome accession G19833, were obtained from the NCBI SRA (BioProject: PRJNA41439); reads were subsampled to obtain 12x coverage of the reference genome accession G19833.

### 2.3 Whole genome assembly and annotation of select accessions

#### 2.3.1 Tissue harvest, library preparation, and sequencing

Four lines, contrasting for three target trait loci [ADP0602 (cv. ‘Sacramento’), ADP0632 (cv. ‘TARS-HT1’), ADP0633 (cv. ‘TARS-HT2’), and ADP0638 (cv. ‘Red Hawk’)], were selected for *de novo* genome assembly. Two seeds per pot were germinated and grown for 17 days, (10 days on light rack then moved to a growth chamber for seven days, 25.5°C day, 15.5°C night, 15 hour day). Leaves were harvested and frozen in liquid nitrogen. High molecular weight DNA was isolated from true leaf tissue from one plant per accession via CTAB extraction and a QIAGEN 500/g Genomic-tip (Qiagen, Germantown, MD) followed by an Amicon buffer exchange (Millipore, Burlington, MA) and filtering of short DNA fragments using the Circulomics Short Read Eliminator Kit (Circulomics Inc, Baltimore, MD) (Vaillancourt et al., 2019). Oxford Nanopore gDNA libraries were prepared using the Ligation Sequencing Kit SQK_-_LSK110 (Oxford, UK) with an input of 3 ug of high molecular weight DNA. Two libraries were made and sequenced on two FLO-MIN106 flow cells for each accession on a MinION or MinIT using the most up to date software available at the time (MinKNOW v20.10.3 (MinION), MinKNOW 20.10.6 (MinIT)). Base calling was performed using Guppy v6.0.1 (https://community.nanoporetech.com/downloads) with the following options: --flowcell FLO_-_MIN106 --kit SQK-LSK110 --calib_detect (config file dna_r9.4.1_450bps_hac.cfg).

To obtain higher sequence coverage for subsequent Nanopore read correction, the same KAPA HyperPlus DNA libraries (described previously) were submitted for resequencing to increase read depth. Read quality was assessed using FastQC v0.11.9 (https://www.bioinformatics.babraham.ac.uk/projects/fastqc/) and MultiQC v1.12 (Ewels et al., 2016). Reads were cleaned using Cutadapt v3.7 (Martin, 2011) removing adapter sequence from the 3’ end of both read one and read two (-a/-A) along with the following parameters: -m 100 -q 30 --trim-n --times 2.

#### 2.3.2 Whole genome assembly

Nanopore reads less than 10kb were removed using seqtk (v1.3-r106; https://github.com/lh3/seqtk). Filtered reads were input into the Flye assembler (v2.9-b1786; Kolmogorov et al., 2019) providing an estimated genome size of 587 Mb (-g .587G) and zero polishing iterations (-i 0). Two rounds of polishing were then performed using all passed Nanopore reads and Racon (v1.4.20; https://github.com/isovic/racon). First reads were mapped using minimap2 (v2.24-r1122; Li, 2021) and then alignments used as input into Racon. One round of Medaka (v1.6.1; https://github.com/nanoporetech/medaka) was then run, again using all passed Nanopore reads and the closest base calling model -m r941_min_hac_g507. Finally, Cutadapt cleaned WGS Illumina reads were used as input into Pilon (v1.24; Walker et al., 2014) for one final round of error correction. First, reads were mapped to the assembly using bwa_-_mem2 (v2.2.1; Li and Durbin, 2009; Vasimuddin et al., 2019) and then alignments were sorted and indexed using SAMtools v1.6 (Li et al., 2009). The sorted bam file was then used as input into Pilon. At each step in the assembly and polishing/error correction pipeline statistics were calculated using assembly-stats (https://github.com/sanger-pathogens/assembly-stats) and gene completeness estimated using BUSCO v5.3.2 (Manni et al., 2021) in genome mode with the embryophyta_odb10 lineage database.

#### 2.3.3 Genome annotation

The four assembled genomes and the common bean reference genome assembly (pvul_v2; Pvulgaris_442_v2.0; Goodstein et al., 2012) were annotated by first generating a species-specific repeat library using the pvul_v2 assembly and RepeatModeler (v2.0.3; Flynn et al., 2020). The repeat library was processed with Protex (v1.2; Campbell et al., 2014) to remove protein-coding genes and Viridiplantae repeats from RepBase (v20150807) were then added to create the final custom repeat library. The genome assemblies were then repeat masked with RepeatMasker (v4.1.2-p1; Chen 2004) using the custom repeat library and the following parameters: -s -nolow -no_is -gff. RNA-Seq reads were aligned to the genome assemblies with HISAT2 (v2.2.1; Kim et al., 2019) using the parameters --max-intronlen 5000 --rna-strandness ’RF’ and the alignments sorted with SAMtools sort (v1.16.1; Li et al., 2009; Danecek et al., 2021). Working gene model sets were predicted for each genome assembly with BRAKER2 (v2.1.6; Brůna et al., 2021) using the soft-masked genome assemblies and the respective RNA_-_seq alignments as hints. Functional annotation was assigned by searching the Arabidopsis proteome (TAIR10; Lamesch et al., 2012), Swiss-Prot plant proteins (v081715; Boeckmann et al., 2003), and PFAM (v35; El-Gebali et al., 2019) and assigning function from the first hit encountered. Metrics to estimate genome assembly and annotation completeness were generated with BUSCO (v5.4.3; Manni et al., 2021) with the embryophyta_odb10 database. Syntenic blocks were identified between the five genomes using GENESPACE (v1.3.1; Lovell et al. 2022). The riparian plot indicating synteny was generated using pyGenomeViz (v1.0.0; https://moshi4.github.io/pyGenomeViz/).

### 2.4 Single nucleotide polymorphism bioinformatics pipeline

Raw reads were quality trimmed using Trimmomatic (v.0.32; Bolger et al. 2014) with the following parameters: LEADING:10, TRAILING:10, SLIDINGWINDOW:4:15, and MINLEN:40. FastQC and MultiQC were used to verify that low quality reads and base-calls were trimmed successfully. Both paired and unpaired reads were aligned to the *P. vulgaris* reference genome (v2.1; https://phytozome-next.jgi.doe.gov/info/Pvulgaris_v2_1) using the Burrows-Wheeler Alignment Tool BWA-MEM (v.0.7.16a; Li 2013). BWA-MEM was run using default parameters, and the options ‘-M’ and ‘-R’ were used for PICARD compatibility and to track sequence read groups. PICARD tools (v.2.20.8; https://broadinstitute.github.io/picard/) and SAMTools (v.1.9; Li et al. 2009) were used to sort the paired and unpaired alignment files, mark duplicates, index alignments, and to collect alignment summary metrics. With the exception of G19833, greater than 95% of the reads from each genotype aligned to the reference genome; for G19833, a lower percentage of reads (93%) aligned to the reference genome due to lower read quality and shorter read length from an older generation Illumina sequencing platform. Overall, the percent of duplicate reads ranged from 7% to 15% across all lines sequenced.

Variant discovery was performed using the Genome Analysis Toolkit (v.4.1.2.0; GATK; McKenna et al. 2010), according to the best practices for ‘SNP and Indel discovery in germline DNA’. With the exception that base quality scores could not be recalibrated due to the lack of a high-quality reference SNP set for *P. vulgaris*. Haplotype calling was performed using the GATK HaplotypeCaller using standard parameters for single-sample GVCF calling. A GATK genomic database was built using GenomicsDBImport with standard parameters and a batch-size of 25. All 242 samples, including the reference genome accession, were included in the genomic database. Lastly, population-level joint-SNP calling was done using GenotypeGVCFs with standard parameters.

SNP filtering was done in several successive stages using GATK, VCFTools (Danecek et al. 2011), and PLINK (Purcell et al. 2007). Prior to filtering, 17,126,881 total variants were identified (Table 1). Initial filtration of biallelic SNPs employed GATK SelectVariants using the functions: --select-type-to-include SNP and --restrict-alleles-to BIALLELIC; a total of 13,356,484 biallelic SNPs remained (Table 1). Then a set of hard-filters were applied using GATK VariantFiltration and the following parameters: -filter “QD < 2.0”, -filter “QUAL < 30.0”, -filter “SOR > 3.0”, -filter “FS > 60”, -filter “MQ < 40.0”, -filter “MQRankSum < -12.5”, and -filter “ReadPosRankSum < -8.0”. After excluding hard-filtered SNPs, a total of 10,586,323 remained (Table 1).

**Table 1.** Single Nucleotide Polymorphism (SNP) filtering and variant counts.

| Stage | Number of Variants |
| --- | --- |
| Raw | 17,126,881 |
| Biallelic SNPs | 13,356,484 |
| GATK hard-filters | 10,586,323 |
| VCFTools filtered (maf, miss, dp) | 2,009,889 |
| PLINK filtered (heterozygosity) | 1,540,725 |
| PLINK filtered (LD-prune) | 206,801 |

To verify the genetic identity of lines included in this study, whole-genome SNP data was compared to SNP-chip data from a previous ADP genotyping study (Cichy et al. 2015). A total of 43 lines with less than 95% concordance between whole-genome and SNP-chip data were excluded from further analysis. Additionally, to reduce the effects of population structure on the GWAS, an additional 16 lines of Meso-American descent were excluded. In total, 183 lines remained for subsequent SNP filtering and GWAS analysis (Supplemental Table S1). To filter SNPs for minor allele frequency, missing data, and read depth, VCFTools v.0.1.16 was used with the following parameters: --maf 0.05, --max-missing 0.9, --min-meanDP 3, --max-meanDP 30 (Danecek et al. 2011). The max-meanDP threshold of 30 was determined using the mean read depth plus two standards of deviation. Following VCFTools filtering, 2,009,889 SNPs remained (Table 1). As VCFTools functions did not include an option for filtering SNPs with high heterozygosity, the vcf file was converted to a plink SNP format for additional filtering using PLINK (v.1.90; Purcell et al. 2007). After creating *.bed, *.ped, and *.map files, the function -- hardy was used to obtain the observed heterozygosity and to generate an *.hwe file. The *.hwe file was parsed using RStudio v.1.3.1056 and R/tidyverse, and a list of SNPs with greater than 10% heterozygosity was generated. The plink option –exclude was used to remove all SNPs with >10% heterozygosity. In total, 1,540,725 SNPs remained after filtering heterozygous SNPs (Table 1). Lastly, the plink2 function –indep-pairwise was used for LD-pruning (https://www.cog-genomics.org/plink/2.0/). A window size of 200kb, a step distance of 1, and an r^2^ threshold of 0.99 was used to remove SNPs in perfect or near-perfect linkage disequilibrium. Due to the inbred nature and high LD of domesticated *P. vulgaris*, a total of 206,801 SNPs remained for subsequent GWAS analysis (Table 1).

### 2.5 *K*-mer GWAS bioinformatics pipeline

After quality trimming raw reads, FastUniq was used to remove duplicate reads (v.1.1; Xu et al. 2012). On average, 7.14% of the reads were identified as duplicates and removed from each sequence file. The *k*-mer GWAS package release v.0.2-beta was used along with the following additional software: R v.3.5.3, python v.2.7.13, KMC v.3.1.0, and GEMMA v.0.96 (Kokot et al. 2017, Zhou and Stephens 2012). Canonical *k*-mers are *k*-mers in which both the forward and reverse complement of the sequence are treated as identical. Canonized *k*-mer counting was run in-parallel on each individual with the following options: -t2, -k31, and -ci2. The -k option specifies the *k*-mer length, and 31 nt *k*-mer length was selected to maximize sequence specificity. The -ci option is the threshold for counting *k*-mers. By setting the threshold to “2”, it means that a *k*-mer must appear at least twice to be counted – which is a reasonable expectation given ∼10x sequence coverage. Then, *k*-mer counting was done a second time with no canonization, and the options -b and -ci0 were used as directed by the *k*-mer GWAS manual. For the canonized counts, approximately 400 million *k*-mers were identified from each individual (Table 2). For the non-canonized counts, approximately 800 million to 1 billion *k*_-_mers were identified from each individual (Table 2). Once canonized and non-canonized *k*-mer counting was complete, the function “kmers_add_strand_information” was used to combine the information from the two counts into a single list of *k*-mers for each individual.

**Table 2.**
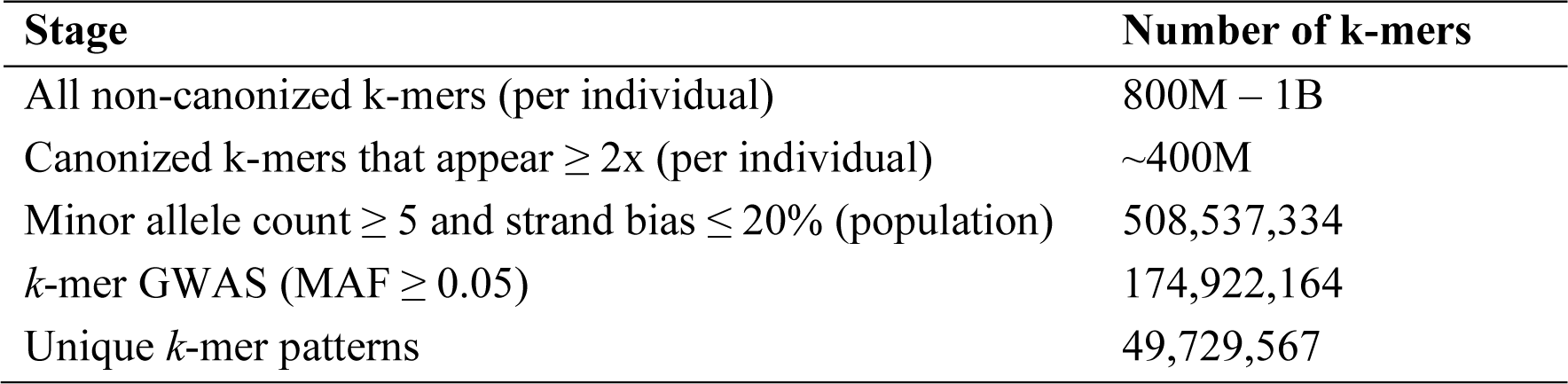
*k*-mer filtering and *k*-mer counts.

The complete list of *k*-mers to include for GWAS was determined based on population_-_level parameters. The same 183 individuals were included with the options –mac 5 and -p 0.2 to compile the complete *k*-mer list. The –mac option determines the minor allele count, or the minimum allowed appearance of a given *k*-mer across the population. The -p option determines the minimum percent of appearance in each strand form, which thereby eliminates *k*-mers with elevated strand-bias. In total, 508,537,334 *k*-mers were identified for GWAS (Table 2). A second list of *k*-mers was generated in the same way after including bean lines with Middle American descent (*n* = 199), and a total of 599,032,584 *k*-mers were identified. In this way, the *k*-mer distributions of “Andean only” and “Andean plus Middle American” could be compared.

Lastly, the *k*-mers presence/absence matrix could be prepared using the function “build_kmers_table”, and emma kinship could be calculated using the function “emma_kinship_kmers” with the option --maf 0.05. After filtering the *k*-mer table for minor allele frequency > 0.05, a total of 174,922,164 *k*-mers remained (Table 2). GWAS analysis was done using the python program “kmers_gwas.py” according to the manual and using the options –kmers_number 100000 and –pattern_counter. The –kmer_number option determines how many *k*-mers to filter from the first step. After testing various thresholds, 100,000 *k*-mers were consistently sufficient for all *k*-mer trait associations. The –pattern_counter option was useful for counting the number of unique presence/absence *k*-mer patterns, and in this dataset, 49,729,567 *k*-mers had unique presence/absence patterns (Table 2). Neighbor joining tree figures were developed using Interactive Tree of Life (https://itol.embl.de/; Letunic and Bork 2021).

### 2.6 Genome-wide association study data analyses

To maintain consistency between SNP-GWAS and *k*-mer-GWAS, GEMMA v.0.96 and the same parameters were used for both analyses. The function emma_kinship was used to calculate kinship, and the phenotypic data was formatted as a *.fam file. A linear mixed model (LMM) which included the kinship matrix was used for all genomic trait associations, and a likelihood ratio test was used to calculate *P* values.

To map *k*-mers associated with each phenotype, Vmatch (v.2.3.0; http://www.vmatch.de/) was used to identify perfect matches to the reference genome. Initially, all associated *k*-mers (exceeding the 5% significance threshold) were parsed from the GWAS results files, and converted to fasta files and vmatch was used with the following options: -p, -d, - complete, and -showdesc 100. The -p option computes and matches the palindrome of each query sequence, while the -d option identifies direct matches for each query sequence. In this way, the *k*-mer strand is accounted for when identifying matches to the reference genome. The - complete option specifies that query sequences must match completely. The -showdesc 100 option, was used as a quick validation of vmatch results. Then the results from vmatch were piped into a results file, and a new *k*-mer GWAS file was reconstructed for significant *k*-mers with mapped genomic positions. In the case of determinacy associated *k*-mers, four *k*-mers were tied for “the most significant”. Four determinate plant lines (ADP0602, ADP0632, ADP0633, and ADP0638) that contained the four most significant *k*-mers were used to generate a de novo assembly of sequences representing the significant *k*-mers. After pulling the raw sequences, SPAdes (v.3.14.0; Bankevich et al. 2012) was used for *de novo* assembly of an 804 bp contig. Default parameters and the option –careful was used to assemble multiple paired-end sequence files from the four selected individuals.

To capture *k*-mers absent from the reference genome due to structural variation, significant *k*-mers for each phenotype from the four sequenced accessions [ADP0602 (cv. ‘Sacramento’), ADP0632 (cv. ‘TARS-HT1’), ADP0633 (cv. ‘TARS-HT2’), and ADP0638 (cv. ‘Red Hawk’)] and the reference genome were converted to FASTA and mapped to their respective genome assembly using Vmatch (v2.3.1) with the parameters -complete -d -p - showdesc 0 -noevalue -noscore. The Vmatch alignment files were then converted to BED format.

### 2.6 Data Availability

Whole genome sequence reads have been deposited in the National Center for Biotechnology Information under BioProject PRJNA1112458 (to be released upon publication). Large files (GWAS results, significant *k*-mers, *de novo* genome assembly and annotations) are available in Figshare under doi (to be made public upon publication).

## 3. RESULTS AND DISCUSSION

### 3.1 Population Structure

In this study, the same WGS dataset was used to produce two marker matrices for association analysis, one composed of SNPs (Table 1) and the other of *k*-mers (Table 2). Two parallel bioinformatic pipelines were run to process Illumina sequence reads into a WGS-SNP matrix using GATK, VCFTools, and PLINK, and into a k-mer matrix using the package released by Voichek and Weigel (2020). In total, this study began with 242 lines including the reference genome accession G19833. To verify the identity of ADP members, historic SNP-chip data (Zuiderveen et al. 2016) was compared to WGS-SNPs and 43 lines with less than 95% concordance were excluded from further analysis. Following GATK hard-filtering of SNPs, a principal component analysis using 100,000 randomly selected SNPs from the remaining 199 lines was performed (Figure 1). In total, 16 lines were identified with Middle American descent and were excluded from downstream association analysis to reduce the effect of population structure. Similarly, *k*-mer counting was performed on the same two population cohorts, one with only Andean members (*n* = 183), and the other with both Andean and Middle American members (*n* = 199). Interestingly, when *k*-mer allele counts were plotted for each cohort, the distributions revealed a core and a unique fraction of the *k*-mers that are specific to Middle American lines (Figure 2).

**Figure 1.**
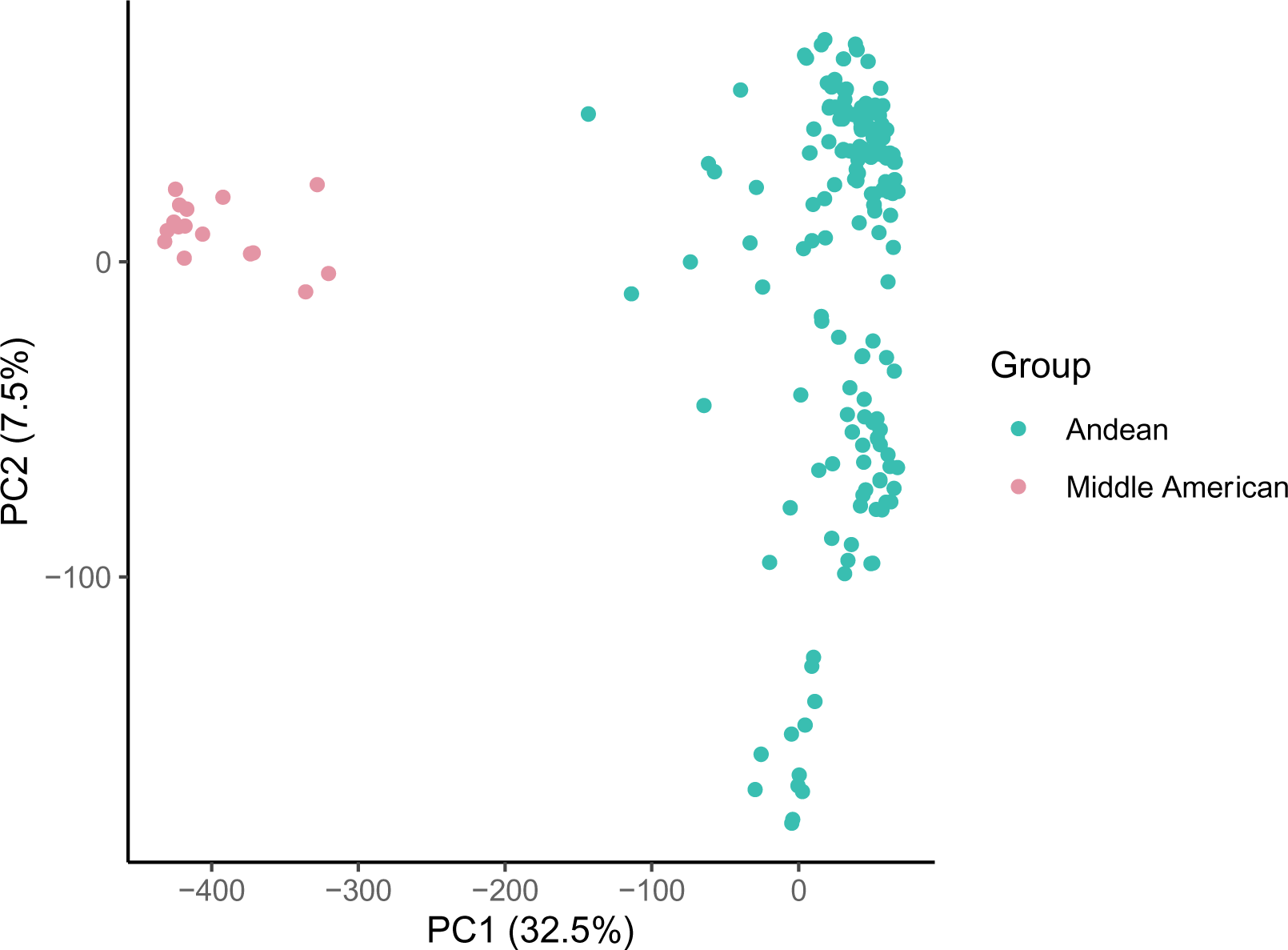
Principal component analysis of single nucleotide polymorphism data differentiating Andean (teal) and Middle American (pink) members of the sequenced panel of lines.

**Figure 2.**
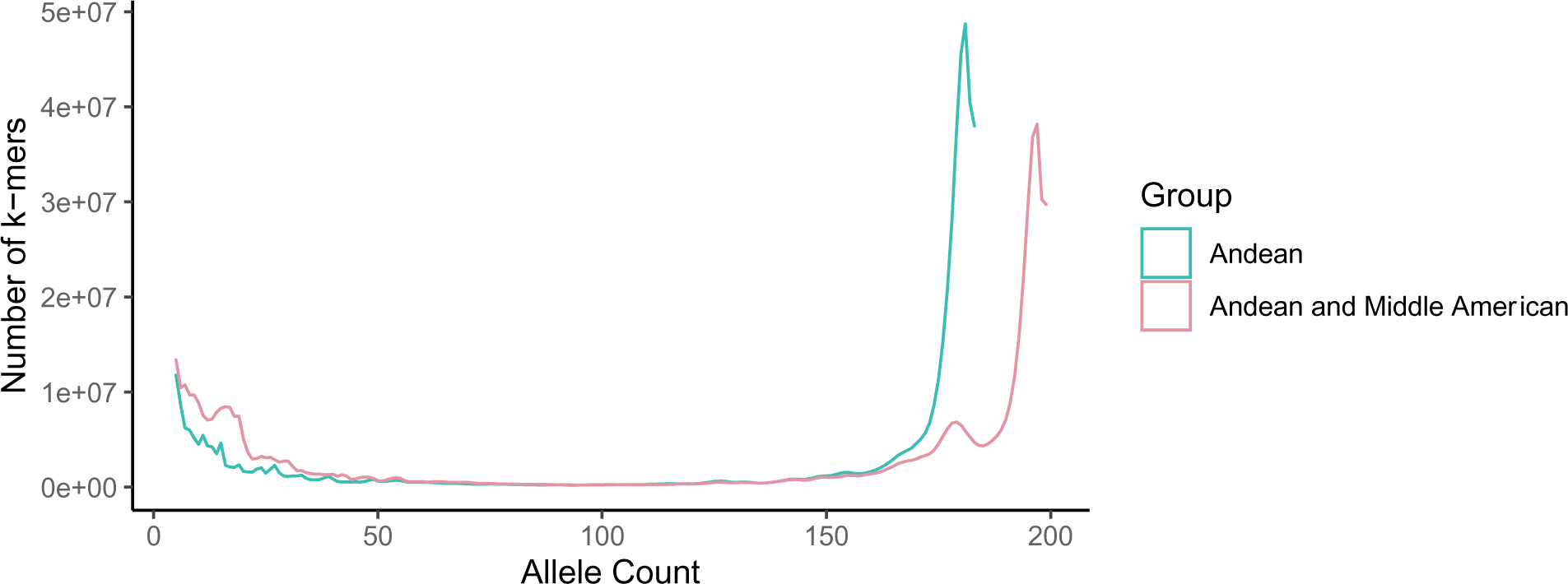
Allele counts for Andean (teal) and Andean and Middle American (pink) *k*-mers. Histogram of *k*-mer allele counts, where core *k*-mers are present in most lines (right), dispensable *k*-mers are present in subsets of the population (middle), and unique *k*-mers are only present in a few lines (left).

### 3.2 Bean Determinacy

As a proof-of-concept, plant determinacy was evaluated as it is a well characterized trait controlled by a single dominant gene, *PvTFL1y*, located on chromosome 1 of the *P. vulgaris* genome (Kwak et al. 2012). The dominant expression of *PvTFL1y* is an indeterminant growth habit with a visible vine, whereas the homozygous recessive expression is a determinate growth habit with terminal flowering (colloquially referred to as a bush-type bean). WGS-derived SNP and *k*-mer association analysis was performed in GEMMA using a Linear Mixed Model that included a kinship matrix produced from each respective marker set. While the SNP markers had predetermined reference genome positions, the reference genome positions of the *k*-mers were not known prior to association analysis, thus, *k*-mer GWAS can be considered a reference-free approach. Initial *k*-mer association analysis only assigned a *P* value to each *k*-mer which exceeded the 5% family-wise error-rate threshold. Of the 42,791 *k*-mers that exceeded the significance threshold for plant determinacy, a total of 20,099 (47.0%) could be directly mapped to the reference genome using Vmatch (Table 3). As expected, highly significant SNPs and *k*_-_mers were identified in the region surrounding *PvTFL1y* on Chromosome 1, and there was exceptional concordance between the SNP and *k*-mer results (Figure 3). Some of the *k*-mers with the most significant *P* values could not be mapped to the reference genome. To determine the genomic context of the most significant *k*-mers, the source reads of the four *k*-mers with the lowest *P* value were retrieved from four determinate lines (ADP0602, ADP0632, ADP0633, ADP0638) that were positive for those *k*-mers and a *de novo* assembly was performed producing a single 804 bp contig (Supplemental Figure S1) that encoded a retrotransposon which has previously been shown to be inserted in *PvTFL1y* (Figure 3, Kwak et al. 2012).

**Figure 3.**
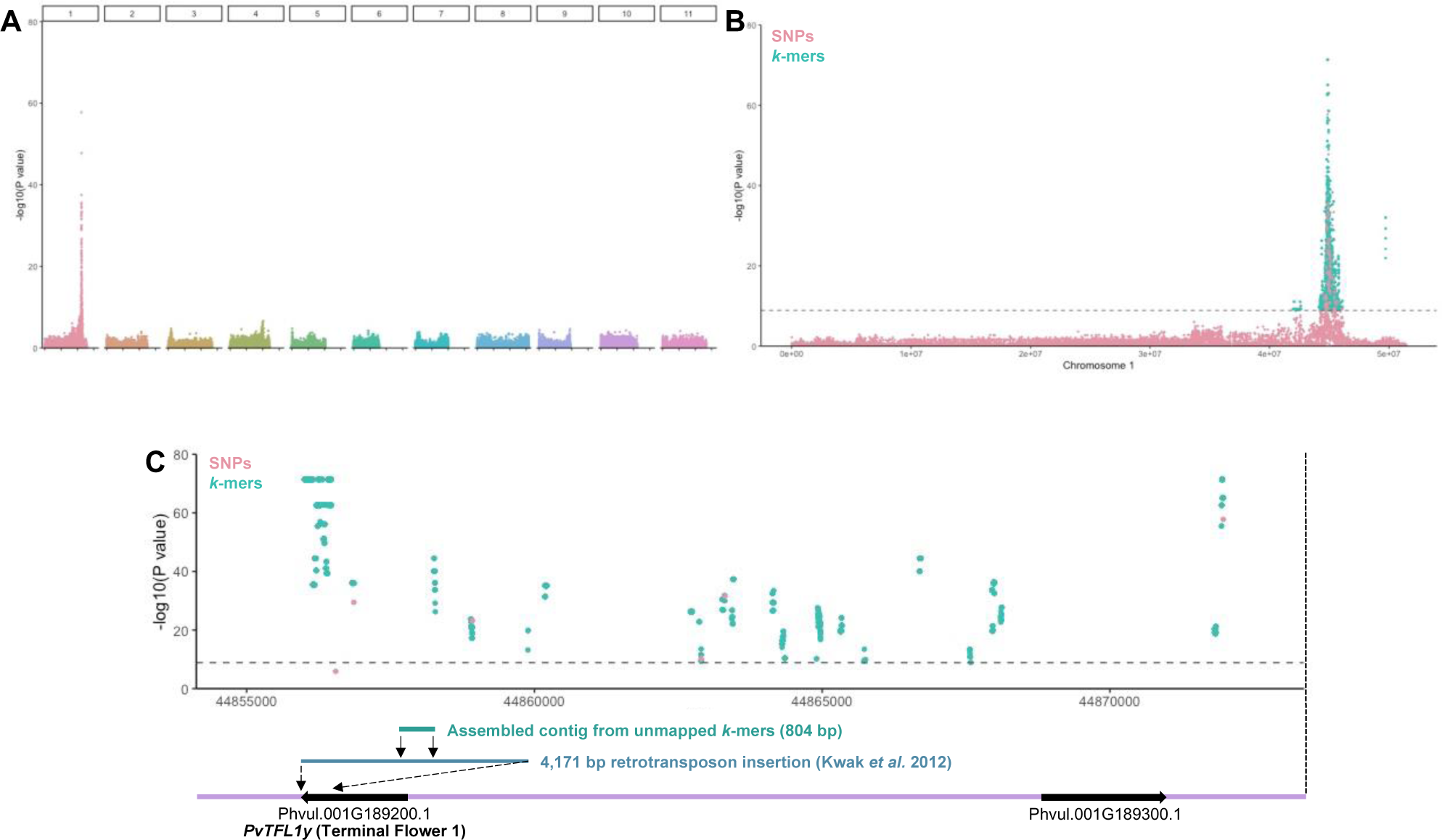
Identification of single nucleotide polymorphisms and *k*-mers associated with plant determinacy. A) Genome-wide SNPs associated with plant determinacy. B) SNPs (pink) and *k*-mers (teal) associated with plant determinacy on Chromosome 1. C) Zoomed-in view of the most significant SNPs (pink) and *k*-mers (teal) relative to local gene positions, including *PvTFL1y* (Terminal Flower 1). An 804 bp contig (teal) assembled from the most significant and unmapped *k*-mers could be mapped directly to the 4,171 bp retrotransposon insertion (blue) identified by Kwak *et al*. (2012).

**Table 3.**
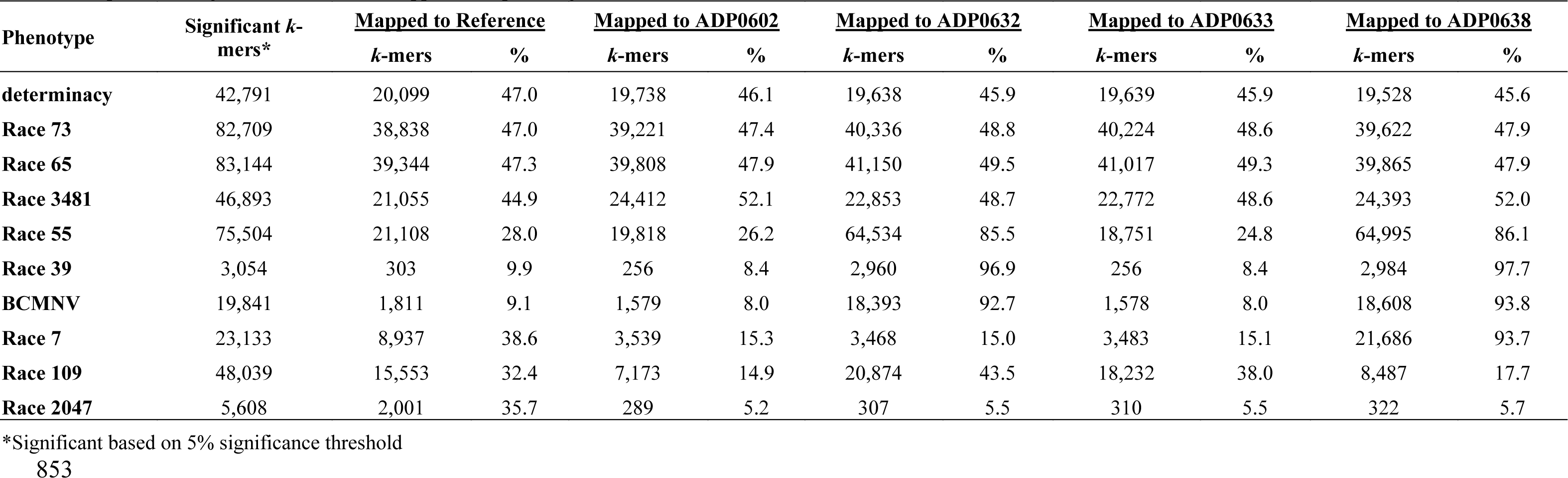
Proportion of significant *k-mers* that mapped to respective genome assemblies.

In the case of bean determinacy, two factors prevented SNP-based GWAS from identifying the causal variant. First, the reference genome accession, G19833, has an indeterminate plant type and therefore lacks the retrotransposon insertion in *PvTFL1y*. Therefore, while the retrotransposon insertion exists within the pan-genome of Andean bean cultivars, its absence from the reference genome prevents discovery using a SNP-based GWAS method that is reliant on a reference genome. Second, SNP-based GWAS is not an effective method for marking structural variants such as insertions. *K*-mer GWAS overcomes both limitations as it is not reliant on a reference genome, and it can identify multiple types of genetic variants— including insertions.

### 3.3 GWAS for disease resistance

In 2016, Zuiderveen et al. published GWAS results for anthracnose resistance on the ADP population using the Illumina BARCBean6K_3 BeadChip with 5,398 SNPs. Using both WGS-SNPs and *k*-mers, we were able to corroborate the results of Zuiderveen et al. (Figure 4, Supplemental Figure S2), identifying marker trait associations (MTAs) in the same three genomic intervals: Pv01 (*Co-1* locus), Pv02 (*Co-u* locus), and Pv04 (*Co-3* locus). In addition to having much higher marker resolution in our study, we also identified significant MTAs for anthracnose Race 2047 on Pv04 – which could not be identified using SNP-chip data. One observation that was immediately apparent from WGS-SNPs and *k*-mers was the relative size of linkage blocks containing the most significant markers. While the MTAs for the *Co-1* locus formed a tight “pyramid-shaped” peak that centered around a central point (Pv01:49,625,120), MTAs for *Co-u* identified a large ∼800 Kb linkage block (Pv02:48,700,000 to 49,500,000) and MTAs for *Co-3* identified an even larger ∼2 Mb linkage block (Pv04: 250,000 to 2,250,000) (Figure 4). Given the size of these linkage blocks, it is difficult to resolve the exact position of potentially causal variants.

**Figure 4.**
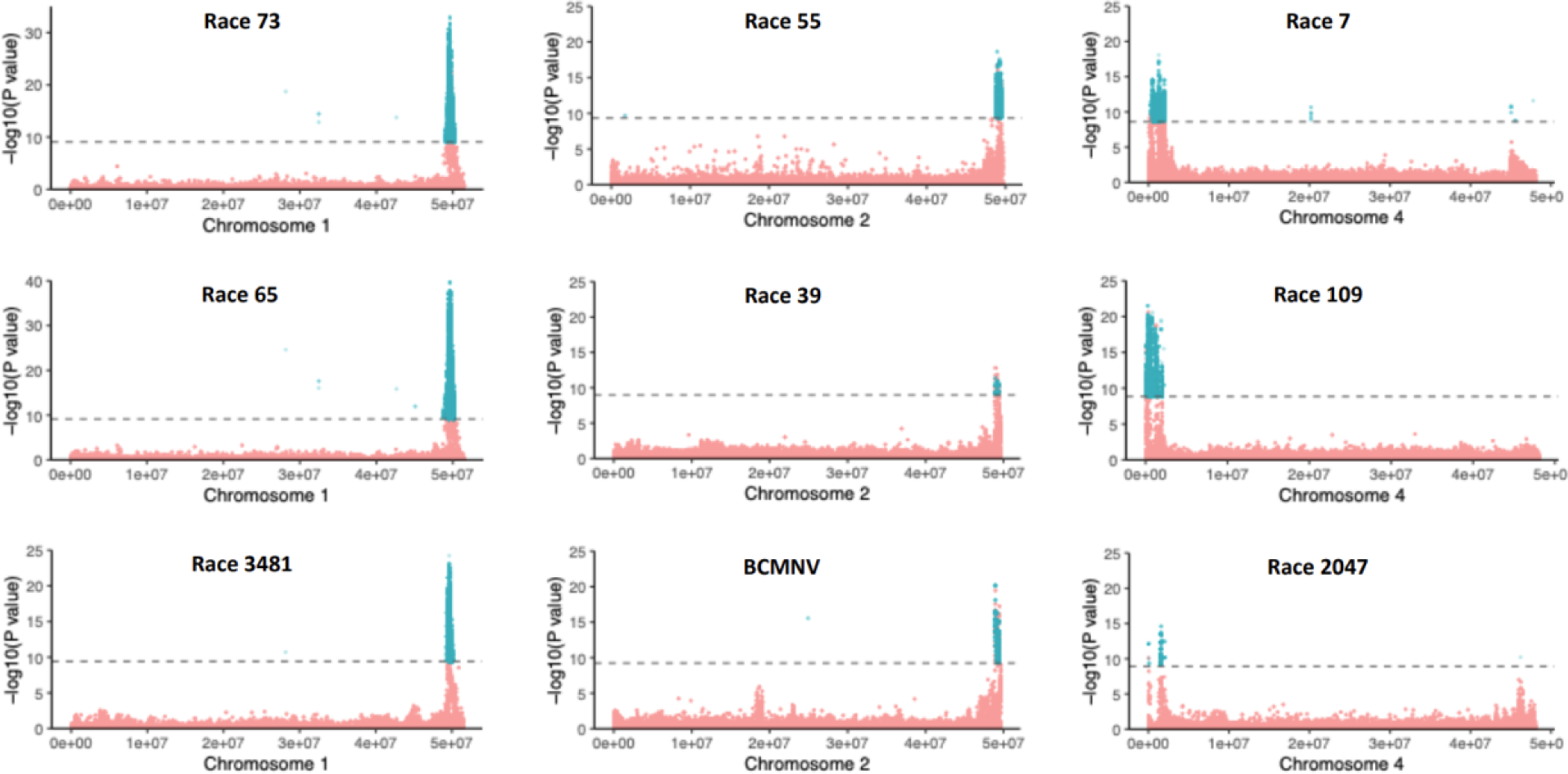
Single nucleotide polymorphisms (SNPs, pink) and *k*-mers (teal) associated with disease resistance phenotypes. The most significant associations for Races 73, 65, and 3481 were identified on Chromosome 1, the most significant associations for Races 55, 39, and BCMNV were identified on Chromosome 2, and the most significant associations for Races 7, 109, and 2047 were identified on Chromosome 4. In every case, mapped *k*-mer associations mirrored the SNP-based results.

The WGS-SNP and *k*-mer GWAS results largely mirrored each other (Figure 4). Similar to plant determinacy, less than half of the significant *k*-mers could be mapped to the reference genome in our disease resistance analyses, and in the case of BCMV resistance (the *I*-gene locus) only 9.1% of the significant *k*-mers could be mapped (Table 3). The inability to map a large proportion of *k*-mers was due to two factors: mapping criteria and *k*-mer absence from the reference genome due to structural variation between accessions. The mapping criteria required that each 31 bp *k*-mer have a perfect match to a unique location in the genome (no multimapping) to reduce false associations. Presence/absence structural variation is well known in plant genomes and certainly, some of the unmapped *k*-mers are *cis*-associated *k*-mers that exist within the pan-genome of Andean beans but are absent from the reference genome accession and potentially represent genetic variability that could underpin phenotypic variation that is absent from the reference genome accession as exemplified with the determinacy locus.

We then generated *de novo* genome assemblies for four ADP accessions with contrasting disease and determinacy phenotypes (Table 4). Each genome assembly was highly contiguous with a minimum contig N50 length of 15 Mb and total assembly size greater than 536 Mb (Supplemental Table S3). Completeness of the genome assemblies revealed >99% complete BUSCO orthologs consistent with a high quality assembly (Supplemental Table S4). Each genome along with the reference genome were annotated for gene models using an identical pipeline to permit comparison of genes across these five genomes (Supplemental Table S5); BUSCO analysis revealed greater than 96% capture of complete BUSCOs (Supplemental Table S4). Alignment of the top 5% significant *k*-mers to the four genome assemblies increased the percent alignable *k*-mers revealing the presence of novel sequences in these genomes (Table 3). Additionally, a much higher proportion of *k*-mers associated with resistance to Race 7 aligned to ADP0638—a line resistant to Race 7.

**Table 4.** Presence and absence of the most significant *k-mer* by trait for lines selected for resequencing and genome assembly.

| **Locus (Chr)** | **Trait** | **G19833\*** | **ADP0632** | **ADP0633** | **ADP0602** | **ADP0638** |
| --- | --- | --- | --- | --- | --- | --- |
| ***TFL1y* (Pv01)** | determinacy | - | + | + | + | + |
| ***I* gene (Pv02)** | BCMNV | - | + | - | - | + |
| ***Co-u* (Pv02)** | Race 55 | - | + | - | - | + |
| ***Co-u* (Pv02)** | Race 39 | - | + | - | - | + |
| ***Co-1* (Pv01)** | Race 73 | - | - | - | + | + |
| ***Co-1* (Pv01)** | Race 65 | - | - | - | + | + |
| ***Co-1* (Pv01)** | Race 3481 | - | - | - | + | + |
| ***Co-3* (Pv04)** | Race 7 | - | - | - | - | + |
| ***Co-3* (Pv04)** | Race 109 | - | - | - | - | - |
| ***Co-3* (Pv04)** | Race 2047 | - | - | - | - | - |

\*Reference genome accession

ADP0632 and ADP0638 are resistant to Race 39, Race 55, and BCMV whereas the reference, ADP0602, and ADP0633 accessions are susceptible to these pathogens. The percentage of significant *k*-mers greatly increased for all three of these traits in the genome assemblies of ADP0632 and ADP0638 relative to the other three genomes (Table 3). These resistance traits have been mapped to Pv02 and significant *k*-mers were detected on the lower arm of the corresponding contigs in the ADP632 and ADP638 (Figure 5). Syntenic blocks were identified between all five genomes and significant *k*-mers associated with resistance to Race 39, Race 55, and BCMV in Pv02 syntenic blocks revealed gene expansion in ADP0632 and ADP0638 (Figure 6). Specifically, in the reference genome within the block defined by Pvul_v2.02G038230 that encodes a vacuolar transporter and Pvul_v2.02G038430 that encodes a PLATZ transcription factor family protein, there are 11 genes annotated as Toll interleukin-1 receptor nucleotide-binding site, leucine-rich repeat (TIR-NBS-LRR) genes. In contrast, in resistant accessions ADP0632 and ADP0638, there are 37 and 32 TIR-NBS-LRR genes, respectively, while in the susceptible accessions ADP0602 and ADP0633 (Figure 6), there is a single and six TIR-NBS-LRR genes, respectively, suggestive that expansion of TIR-NBS-LRR genes and subsequent mutation resulted in paralog(s) encoding resistance to each of these pathogens.

**Figure 5.**
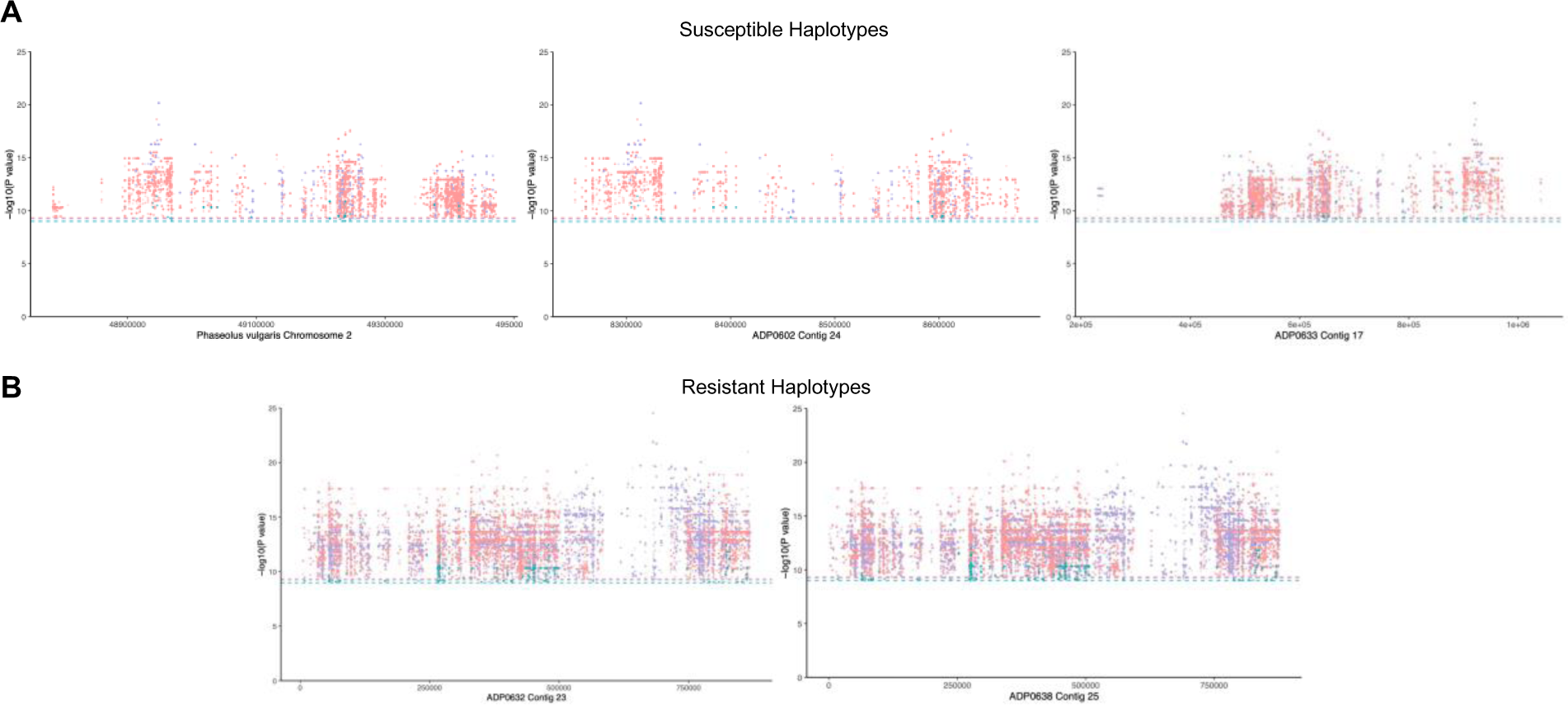
*k-*mers associated with resistance to BCMV and Anthracnose Races 39 and 55 aligned to the reference genome and genome assemblies of ADP0602, ADP0632, ADP0633, and ADP0638. In each case, the chromosome or contig with the most associated *k-*mers is displayed. Race 39 associations (teal), Race 55 associations (pink), and BCMV associations (purple). For each trait, the 5% significance threshold is indicated with a dotted line. A) Susceptible *k*-mer haplotypes and B) resistant *k-*mer haplotypes.

**Figure 6.**
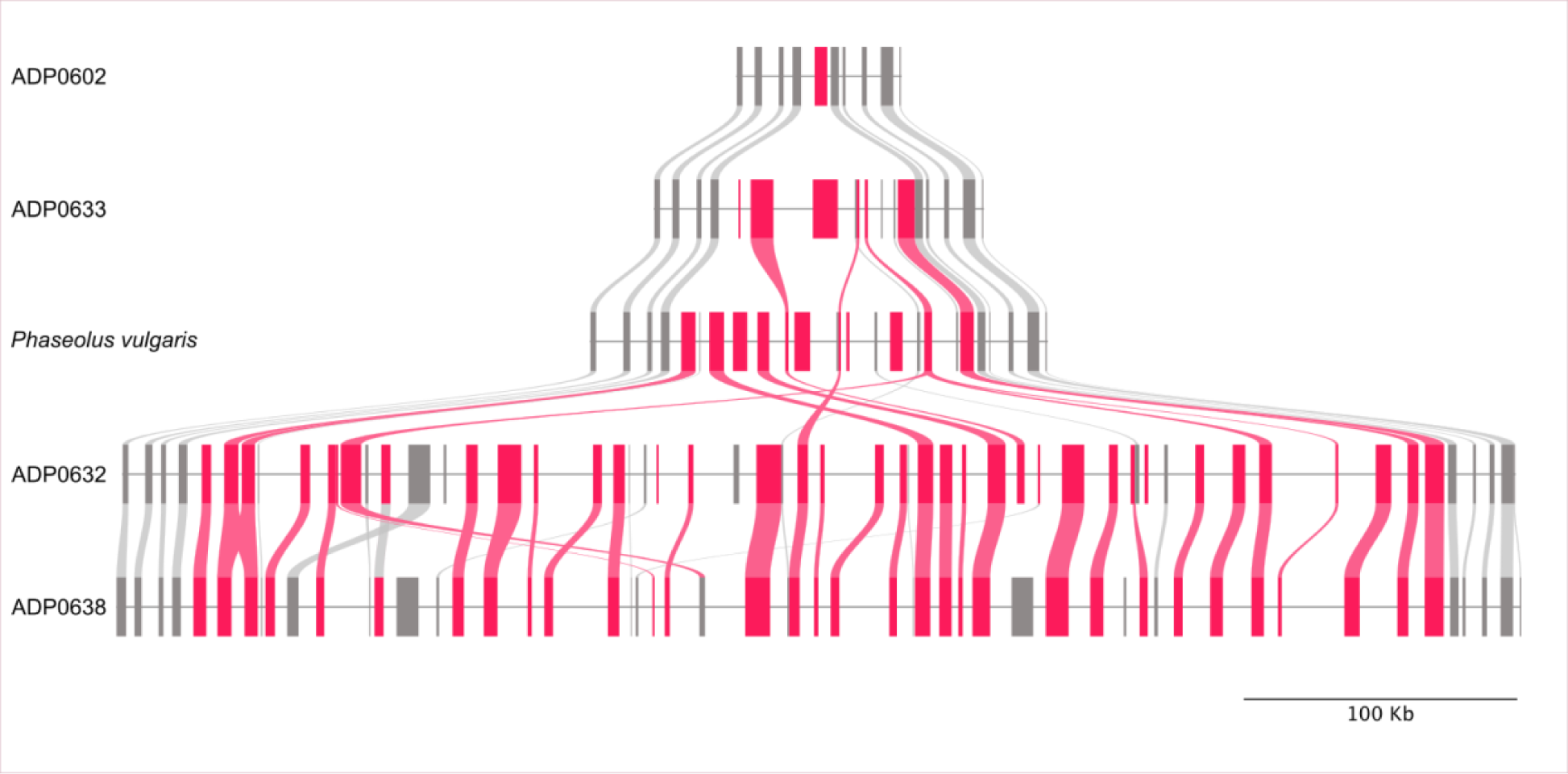
Gene family expansion in Toll interleukin-1 receptor nucleotide-binding site, leucine-rich repeat genes associated with resistance to three pathogens. Riparian plot showing a syntenic region of Chromosome 2 among five common bean genomes (ADP0602, ADP0633, *Phaseolus vulgaris* v2, ADP0632, and ADP0638). Toll interleukin-1 receptor nucleotide-binding site, leucine-rich repeat genes and lines indicating synteny are colored in red; other genes and their synteny are colored in grey.

### 3.4 *k*-mer haplotypes as predictors of disease resistance in the ADP

We constructed a neighbor-joining tree for the 183 ADP lines evaluated in this study and overlaid the phenotypic data and the presence/absence patterns of the most significant *k*-mers in genomic regions of interest: Pv01 MTAs (Supplemental Figure S3), Pv02 MTAs (Figure 7), and Pv04 MTAs (Supplemental Figure S4). This permitted the visualization of the genetic relationships between lines and how specific *k*-mer presence/absence correlates with each phenotype. In the case of Pv01 MTAs, although the phenotypic responses are all correlated, the most significant *k*-mer for Race 73 is associated with resistance whereas the most significant *k*_-_mers for Races 3481 and 65 are associated with susceptibility (Supplemental Figure S3). Furthermore, for Race 2047, it is apparent that only a few individuals are resistant to Race 2047, which also explains the difficulty of mapping a resistance locus for Race 2047 (Supplemental Figure S4). Lastly, based on *k*-mer presence/absence patterns for all traits included in this study, we can predict that the reference genome accession G19833 is susceptible to all the anthracnose races tested and does not have the *I*-gene for BCMV resistance. Unfortunately, G19833 was not included in the ADP, and seed for G19833 was not available when disease phenotyping was being done.

**Figure 7.**
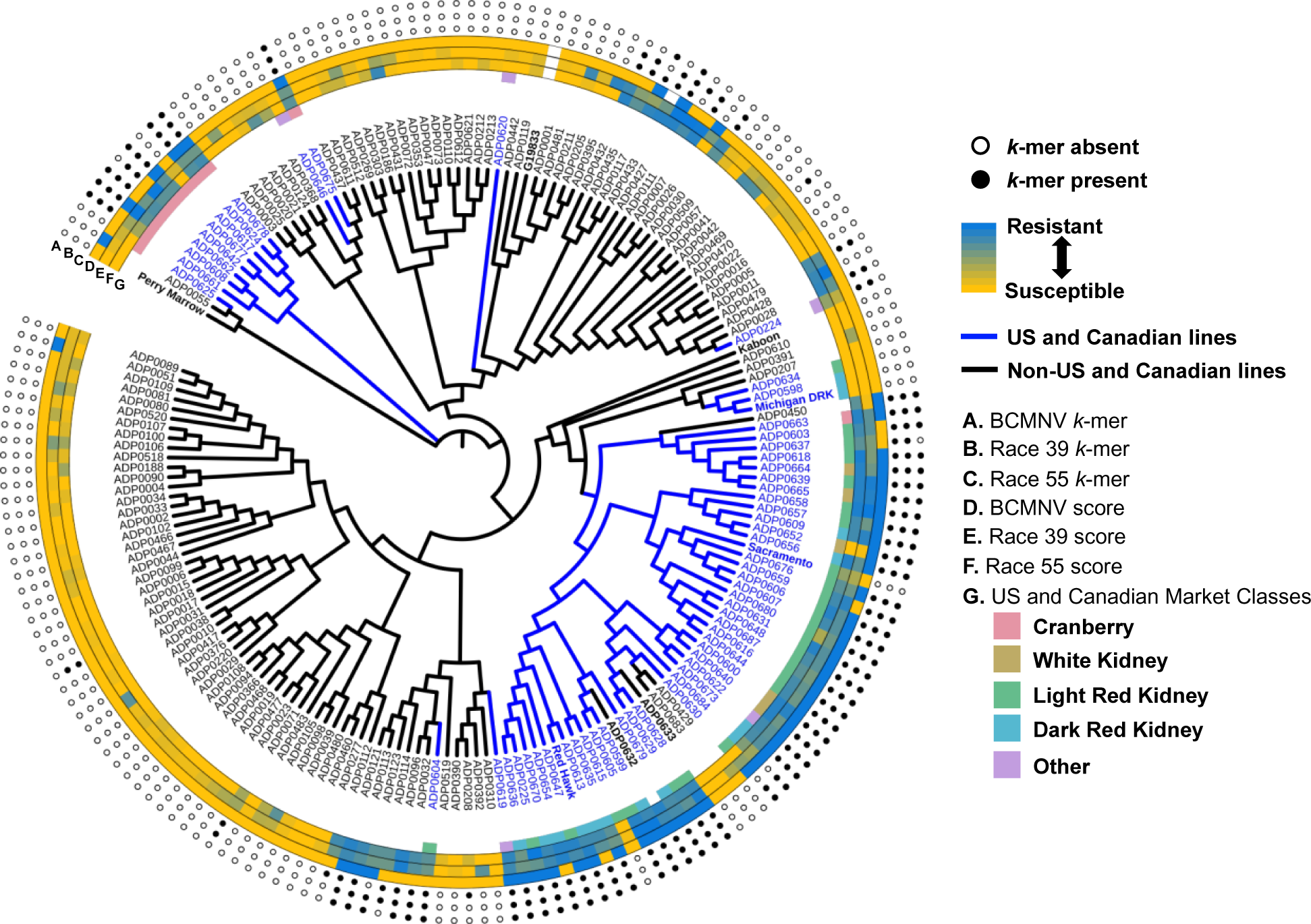
Neighbor joining tree of the Andean diversity panel, with US and Canadian lines (blue branches), chromosome 2 *k*-mer trait associations (A-C), and phenotypic data (D-F). The presence (closed circle) or absence (open circle) of the most significant *k*-mer for each respective trait is displayed in the outer 3 rings (A-C). The phenotypic data for BCMNV (D), Race 39 (E), and Race 55 (F) are displayed on a continuum from susceptible (yellow) to resistant (blue). On the inner-most ring (G), the market classes ‘cranberry’ (pink), ‘white kidney’ (tan), ‘light red kidney’ (green), ‘dark red kidney’ (blue), and ‘other’ (purple) are displayed for the US and Canadian lines.

### 3.5 Genomic evidence for *I*-gene selection in North American breeding programs

The geographic origins of the 183 ADP lines included in this study are diverse and include lines collected from Africa, South America, Central America, North America, the Caribbean, Asia, and Europe. Equally diverse are the seed market classes which can differ in seed coat color and pattern, and seed size. Each region or country has specific preferences that are deeply rooted in cultural traditions, and the names of specific market classes are as variable as the traditions that grow and produce them. The four primary Andean bean market classes in the U.S. and Canada are cranberry beans, white kidneys, light red kidneys, and dark red kidneys (Figure 7). Due to the seed coat pattern and seed shape of cranberry beans, typically cranberry beans are only crossed with other cranberry beans to maintain market class standards.

Alternatively, the three classes of kidney beans are regularly interbred with each other since they primarily differ in seed coat color—which is controlled by only a few major genes (McClean et al. 2002, McClean et al. 2018). Development of a neighbor-joining tree for the ADP lines included in this study supports our knowledge of historical breeding efforts for cranberry and kidney beans in the United States and Canada. As expected, the kidney bean varieties all cluster together on a single branch that is distinct from the Cranberry beans and beans from other parts of the globe.

In addition to breeding for various agronomic characteristics such as seed yield, plant architecture, and harvestability, U.S. and Canadian breeding programs have also emphasized breeding for disease resistance traits. One disease that can severely impact harvest potential and contaminate seed lots is BCMV. In North America, the primary genetic defense against BCMV has been the incorporation of the *I*-gene into advanced breeding lines and commercial varieties. After superimposing Pv02-associated traits onto the neighbor-joining tree, there is clear evidence of selection for the *I*-gene in North American breeding germplasm and varieties. Compared to Andean beans originating from other regions, a much higher proportion of the North American lines are positive for *k*-mers associated with the *I*-gene. Additionally, by selecting for *I*-gene resistance in North American cranberry and kidney beans, it appears that the genetically linked *Co-u* locus for anthracnose resistance was also selected for inadvertently (Figure 7).

## 7 CONCLUSIONS

Low resolution markers such as SNP chip and GBS datasets coupled with high linkage disequilibrium limit in common bean the detection of causal genes and development of accurate molecular markers for breeding. Using the ADP, we demonstrate how access to WGS data permitted high resolution association of variants, i.e., *k*-mers, that can be associated with key phenotypes. We also demonstrate that local expansion of TIR-NBS-LRRs in select accessions are associated with resistance to a viral and fungal pathogen. In this study, not only have markers been identified that can be used in molecular breeding but novel sequences outside the reference genome that confer key agronomic traits have been identified. This study advances our understanding of how the pan-genome contributes to phenotypic variation and further demonstrates the importance of a pan-genome in identifying markers for breeding.

## AUTHOR CONTRIBUTIONS

ATW generated sequence data. BV generated nanopore sequence and performed whole genome assembly. JPH performed genome annotation and analyzed *k*-mers in a genome context. JB performed bioinformatic analyses. HEW supported greenhouse production and helped confirm plant growth habit. EMW provided the ADP seed. All authors contributed to the data analysis. ATW, JDK, and CRB conceived of and designed the study. All authors contributed to manuscript writing, editing, and approval.

## ACKNOWLEDGMENTS

This project was funded by an award from the USDA-NIFA AFRI Education and Workforce Development Program (Award No. 2019-67012-29717) to ATW and funds from USDA-NIFA (2022-67013-37119) to CRB. The authors acknowledge funding from the Michigan State University Plant Resilience Institute, Michigan State University, the University of Georgia, Georgia Seed Development, and the Georgia Research Alliance.

