## Supplementary Figures for "*K*-mer Genome-wide Association Study for Anthracnose and *BCMV* Resistance in the Andean Diversity Panel"

^1^Archer Daniels Midland Company, New Plymouth, ID; ^2^Department of Plant, Soil and Microbial Sciences, Michigan State University, East Lansing, MI; ^3^Plant Resilience Institute, Michigan State University, East Lansing, MI; ^4^Department of Plant Biology, Michigan State University, East Lansing, MI; ^5^Center for Applied Genetic Technologies, University of Georgia, Athens, GA. ^6^Department of Crop and Soil Sciences, ^7^Institute of Plant Breeding, Genetics & Genomics, University of Georgia, Athens, GA.

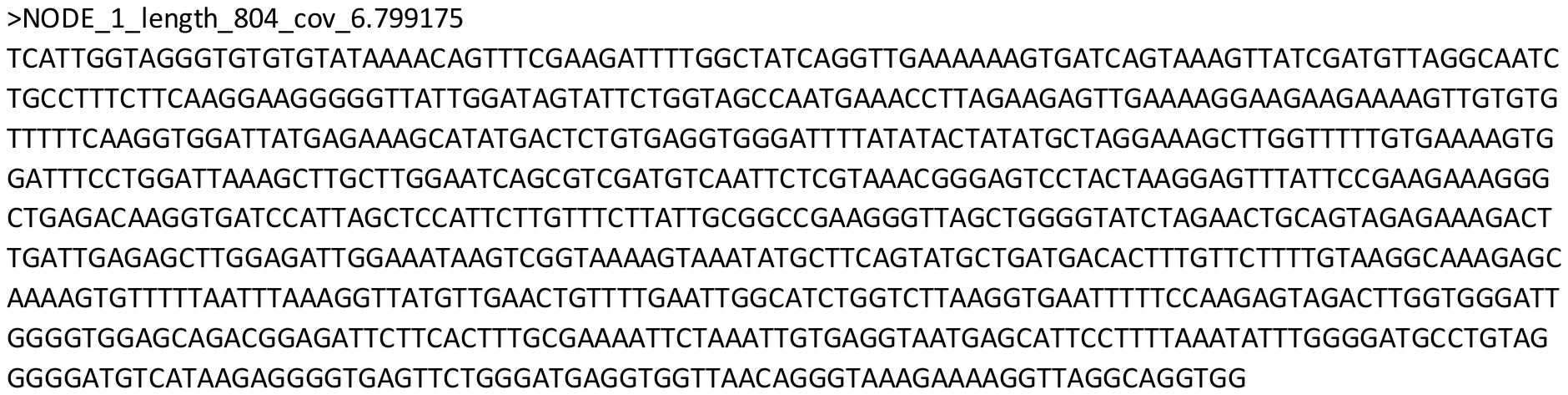

**Supplemental Figure S1: Determinacy contig assembled from sequence reads of unmapped *k*-mers**

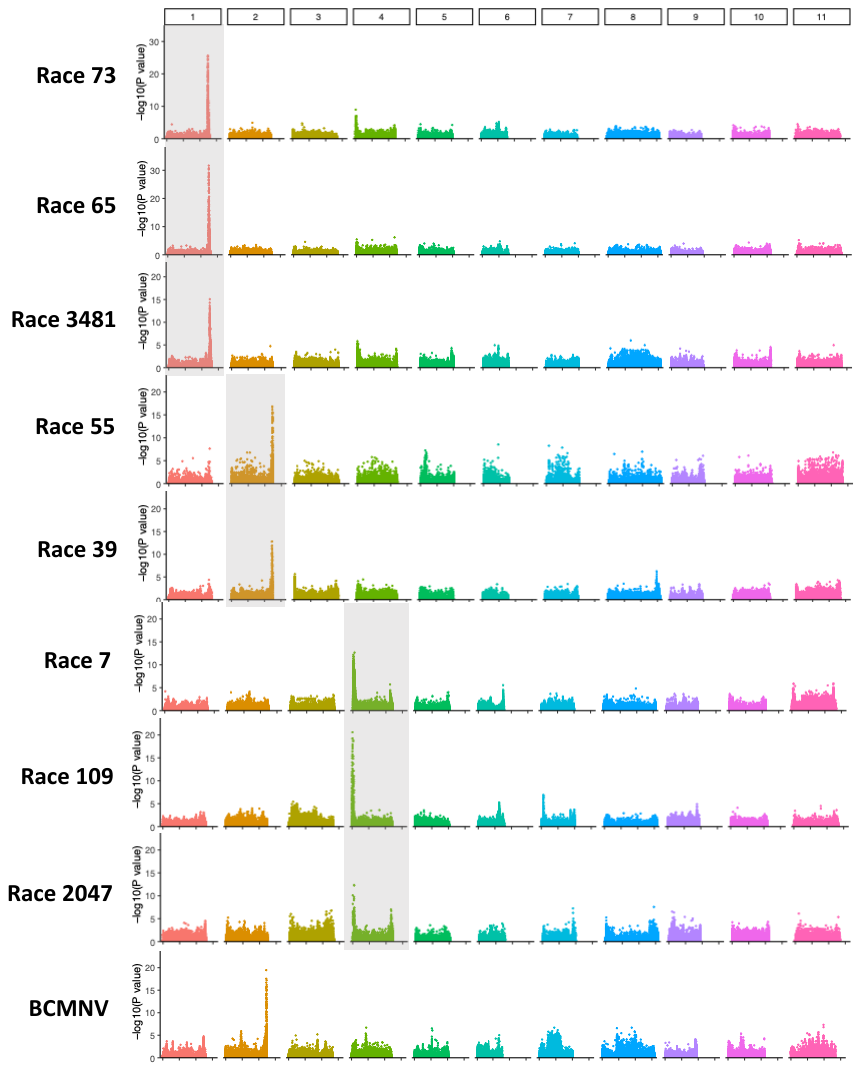

**Supplemental Figure S2: Genome-wide single nucleotide polymorphisms (SNPs) associated with disease resistance phenotypes.**

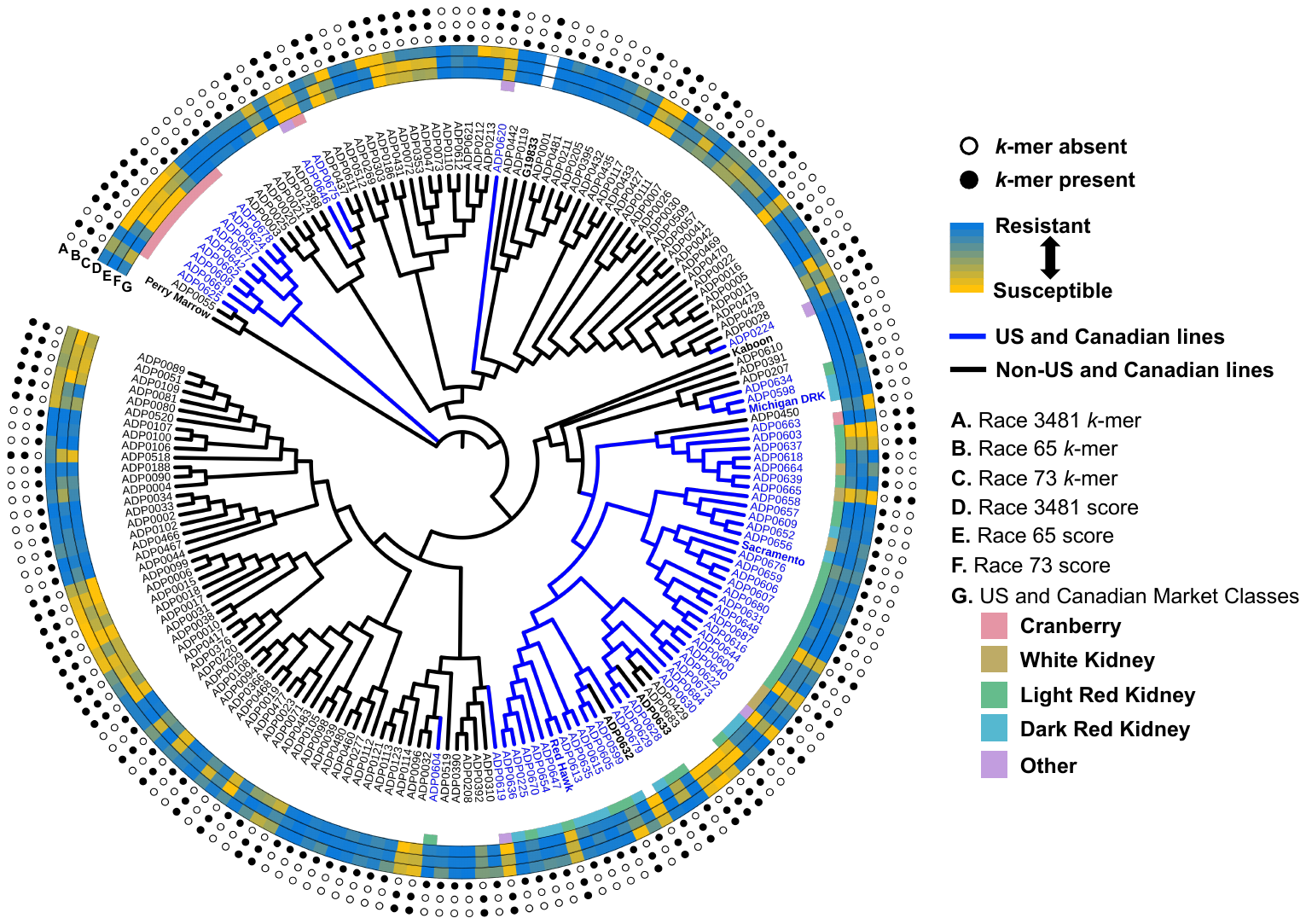

**Supplemental Figure S3: Neighbor Joining Tree and chromosome 1 *k*-mer trait associations.**

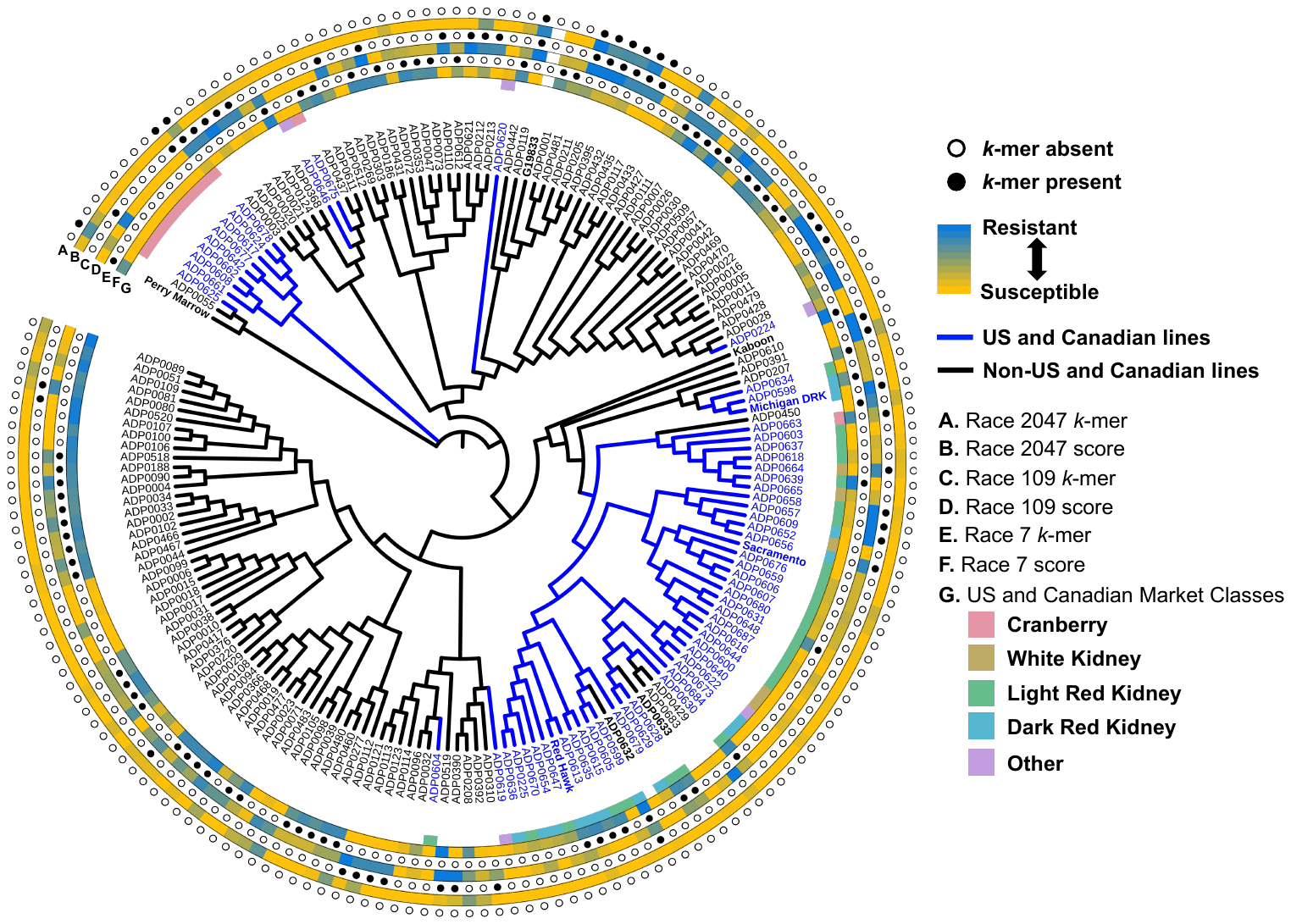

**Supplemental Figure S4: Neighbor Joining Tree and chromosome 4 *k*-mer trait associations.**
